# The role of floral traits in community assembly process at high elevations in Lesser Himalaya

**DOI:** 10.1101/2022.02.26.482103

**Authors:** Mustaqeem Ahmad, Sergey Rosbakh, Solveig Franziska Bucher, Padma Sharma, Sonia Rathee, Sanjay Kr. Uniyal, Daizy R. Batish, Harminder P. Singh

## Abstract

1. Ecological theory postulates that plant trait research should consider multiple traits related to different organs and/or ontogenetic stages as such traits represent different ecological niche axes. Particularly, floral traits have been suggested to play an important role in assembling plant communities along environmental gradients as they determine the reproductive success, one of the key functions in plants. Yet, the predictive power of floral traits in community assembly research remains largely unverified empirically.
2. We analyzed the predictive power of six floral traits of 139 herbaceous species for inferring community assembly process in twenty-one sites located along an elevation gradient in Lesser Himalaya ranging from 2,000 to 4,000 meters above sea level. The floral trait variability along the gradient was analyzed using community-weighted trait mean (CWM) values and functional diversities (FD) calculated for each of the study communities.
3. The CWM values for onset of flowering and flower display area increased significantly with increasing elevation, whereas specific flower area showed an opposite pattern. In combination with convergence in onset of flowering and specific area (i.e., lower FD values in high elevation sites), these patterns suggest that abiotic filtering and plant-pollinator interactions affected the floral trait composition of the communities studied. Increasing low-temperature stress towards high-elevation sites selected for late-flowering species that produce resource-intensive flowers with larger display areas.
4. Low pollinator abundancy and activity in high elevation, could also explain why these traits were selected in the study communities. Delayed flowering with increasing elevations might facilitate the phenological overlap of plants and their pollinators, as pollinator activity at higher elevation peaks in the second half of the vegetation period. The dominance of species with low specific flower area and larger display area in high elevation communities were attributed to the increased flower longevity and attraction of pollinators, respectively, to maximize pollination success under pollinator scarcity.
5. *Synthesis*. Our study provides empirical support of the recent argument that floral traits contribute considerably to the assembly of plant communities along environmental gradients. Thus, such traits should be included into community assembly research agenda as they represent key growth and survival ecological functions.

## Introduction

Understanding the processes shaping the structure and composition of biological communities along environmental gradients, i.e., the community assembly mechanisms (Keddy, 1992), is one of the fundamental aims in ecological research. Disentangling abiotic and biotic factors selecting for or against species from the regional to local species pools allows us not only to understand the current community compositions but can also help to answer the question how these communities will respond to future environmental changes (Götzenberger et al., 2012).

Traditionally, community assembly processes have been scrutinized through the powerful lenses of trait-based approaches (McGill et al., 2006). These methods are built on the paradigm that functional traits, that is the physiological, morphological or phenological characteristics of a plant individual which directly affect its growth, reproduction, and survival (Violle et al., 2007), determine species’ responses to abiotic environment and biotic interactions (Pillar et al., 2021). Therefore, trait variability within a community reflects the outcomes of abiotic filtering and biotic interactions which are two key assembly rules (Bello et al., 2013; Spasojevic & Suding, 2012) across different scales (Kraft et al., 2008, Weiher et al., 2011).

There is a common agreement that, to maximize our understanding of trait-based community assembly, traits related to different organs should be considered as they relate to different ecological niche axes (Grime et al., 1997; Laughlin, 2014). However, the published studies are biased strongly towards traits connected to vegetative growth, survival and resource acquisition (i.e., vegetative traits) with traits related to sexual reproduction being largely ignored (E-Vojtkó et al., 2020; Jiménez□Alfaro et al., 2016; Rosbakh et al., 2018), therefore, including these traits into the current research agenda is important to predict complex environmental effects on trait composition within communities (E-Vojtko et al., 2020; Poschlod et al., 2013; Rosbakh et al., 2018).

In this context, floral traits, physiological, morphological and phenological characteristics of flowers and inflorescences and the way they are pollinated (Klotz et al., 2002), are of a particular interest. Several floral traits, such as the number of flowers, flower display area, scent, and petal colour, determine the success of reproduction, one of the main functions in plants (Barrett, 2010; Soltis & Soltis, 2014). Thus, to maximize plant reproductive fitness, these traits display strong intra- and interspecific variability along environmental gradients reflecting adaptations in plants to specific abiotic conditions and/or pollinator availability (E-Vojtkó et al., 2020). For example, soil moisture availability limits plant investment in flowers with smaller floral area in dry habitats (Lambrecht & Dawson, 2007). Further, Burkle and Irwin (2010) reported that low soil nitrogen content has direct effects on survival and adult fecundity, and results in increased length of flowering stalks. These findings suggest that, at the community level, abiotic filtering could select floral traits and thus leads to shifts in community functional trait composition. Another potential driver of community assembly process in flowering plants is plant-plant and plant-pollinator interactions, for which previous research has indicated that some floral traits (e.g., flower morphology, colour, and scent) can be involved in competition and facilitation via pollinators (Sargent & Ackerly, 2008). Pollinators, particularly, bees can differentiate between flower temperatures which influences their behaviour as they prefer warm flowers (with larger display area) and warm nectar over unheated ones (Dyer et al., 2006). Norgate et al. (2010) reported that this behaviour becomes more pronounced with decreasing temperatures. Learning behaviour of pollinators may indirectly enhance successful pollination via their preference for some colours through their ability to associate flower temperature with specific colours (van der Kooi et al., 2019). Additionally, this learning behaviour can also affect flowering communities through frequency of their visitation as inexperienced pollinators may be attracted to larger display areas whereas experienced pollinators prefer greater floral rewards (Makino & Sakai, 2007).

Finally, some phenological traits, such as onset and duration of flowering, can be affected by both, abiotic filtering, and biotic interactions (Bucher & Römermann, 2020; Ulrich et al., 2020). Temperature has been reported to influence flowering phenology as higher temperatures lead to an advance in first flowering day (Menzel et al., 2006). Similarly, temperature also displays strong association with flowering duration, with warmer temperature leading to extended flowering period (Bucher & Römermann, 2020; Campbell & Halama, 1993). On the other hand, biotic interactions via pollinators also impact flowering duration as animal-pollinated flowers have a higher flowering duration as compared with wind-pollinated plants (Bucher & Römermann, 2020; Rabinowitz et al., 1981).

The aim of this study was to examine the role of six selected floral traits – onset of flowering, flowering duration, flower number, flower investment, flower display area and specific flower area – for inferring community assembly process along an elevational gradient in the Lesser Himalaya. Using *in-situ* measured trait data for 139 herbaceous species occurring in 21 sites ranging from 2,000 to 4,000 m above sea level (a.s.l.), we analysed variation in community-weighted trait mean (CWM) values and functional diversities (FD), two key metrics of community functional structure (Ricotta & Moretti, 2011), to infer the community assembly processes. Specifically, given the increasing low-temperature stress towards higher elevations and high frost-sensitivity of flowers (Wagner et al., 2011), we anticipated a significant shift in plant strategies towards stress-avoidance by delaying the onset of flowering and consequently shorter flowering duration in high elevation communities (Ahmad et al., 2021; Bucher & Römermann, 2020; Table 1). In addition to that, the short growth period with frequent and severe frosts coupled with generally limited soil resources was expected to select for species which allocate their resources to vegetative growth rather than to sexual reproduction (Laiolo & Obeso, 2017; Rosbakh & Poschlod, 2021). In terms of floral traits, we hypothesized that communities at higher elevations are assembled by species which produce fewer flowers with smaller specific flower area and invest less into flower biomass and consequently have smaller floral display areas. In other words, the strong abiotic filtering at high elevations should lead to significant decrease in the CWMs and stronger trait convergence (low FD values) in all six traits (except for increasing CWMs for onset of flowering) with increasing elevation.

**Table 1.** Overview of floral traits studied and their hypothesized role in the community assembly process with increasing elevation.

| **Traits** | **Abiotic filtering** | **Biotic interactions** |
| --- | --- | --- |
| Onset of flowering | Decreases as a stress-avoidance adaptation. Trait convergence as a result of strong environmental filtering | Delayed and synchronized onset of flowering to maximize pollination success.<br>Highest trait values and trait divergence in lower sites due to high competition for pollinators |
| Duration of flowering | Shorter as onset of flowering is comparatively later | Longer to increase chance to be pollinated. Trait divergence in lower elevation due to comparatively high competition for pollinators.<br>Higher elevation: trait divergence due to facilitation effects |
| Flower number | Fewer at higher elevations due to resource limitations and risky sexual reproduction | More at higher elevations (clustering of flowers) for increased pollinator attraction and visitation.<br>Strong trait convergence due to high facilitation effects |
| Specific flower area | Smaller at higher elevations due to low resource availability | Increases along elevation to ensure successful pollination and fertilization. Trait divergence in lower elevation due to competition for pollinators |
| Floral display area | Smaller at higher elevations due to fewer resources and uncertain sexual reproduction | Larger at higher elevation to attract pollinators. Strong trait convergence at higher elevation due to high facilitation effects |
| Flower investment | Decreases with increasing elevation because of increasing unfavorability for sexual reproduction | Higher in higher elevation to attract pollinators and ensure successful sexual reproduction. Strong trait convergence due to high facilitation effects. Trait divergence in lower elevation due to plant-plant competition for available pollinators |

Alternatively, we hypothesized that elevation-specific plant-pollinator relationships also affect functional composition of the communities. Considering that higher elevational sites observe generally sparse vegetation and lower pollinator abundance (Körner, 2021; Trunschke & Stöcklin, 2017), we expected to observe a pattern of delayed and synchronized flowering that continues for a longer period at high-elevational communities as a strategy to maximize pollinator visits and thus to increase the possibility of successful pollination. In other words, the CWMs for the onset of flowering and flower duration were expected to decrease and increase, respectively, along the elevational gradient. The increasing role of facilitation with increasing elevation (i.e., synchronized flowering will attract pollinators and thereby increase the chances of pollination for all community members; Tachiki et al., 2010) should be reflected in the FD variation. Particularly, comparatively higher competition for pollinators in lower elevations was anticipated to result in trait divergence (higher FD at lower than at higher elevation). With respect to the decreasing pollinator abundance along the elevational gradient, we further hypothesized an increase in flower number, specific flower area, flower display area and flower investment in high elevation communities, as it could be advantageous to enhance pollinator attraction in fragmented alpine habitats combined with a short flowering season and generally low pollinator abundance (Fabbro & Körner, 2004; Zhu et al., 2010; Körner, 2021). That is, the CWM and FD values should increase and decrease with elevation, respectively.

## Materials and methods

### Study system

The study was conducted in the Dhauladhar mountain range, Lesser Himalayas (Himachal Pradesh, NW India; Fig. 1). The region is situated at the junction of western, north-western, and trans-Himalayan ranges and is characterized by high levels of floral and faunal diversity due to the combination of diverse geomorphological, geological, and climatic factors (Ahmad, 2021). The study region has a typical alpine relief, with steep mountain peaks reaching up to 6,200 m a.s.l.. The bedrocks of the Dhauladhar ridge are mainly composed of granite and gneiss (Singh & Singh, 1987). The region has a temperate climate with the growth period lasting from April to September in lowlands with average temperatures of 20°C (https://www.adb.org/). There is a decrease in mean annual temperatures with a lapse rate of 6.5° K km^-1^ of elevation (Kattel et al., 2013). The precipitation pattern in Himachal Pradesh is mainly governed by western disturbances and southwest monsoon (Jaswal et al., 2015). Mean annual precipitation in the region is about 2,900 mm a^-1^ (Jaswal et. al., 2014) with a pronounced rainy season from July to September (Singh & Singh, 1987).

**Figure 1.**
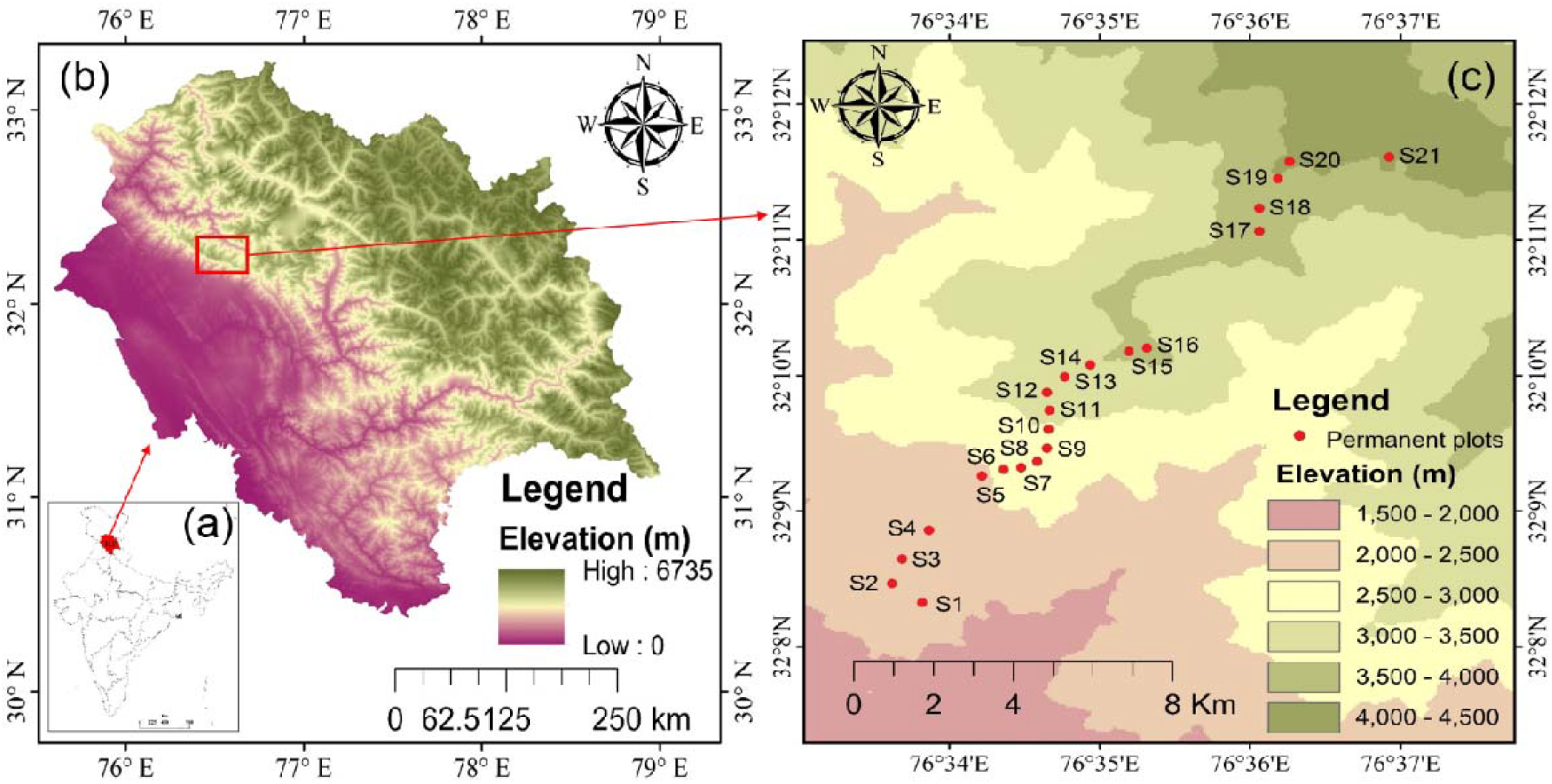
The location of the study sites in India (a) and Himachal Pradesh (b), and their spatial orientation along the elevational gradient (c). See Appendix 2 Table S1 for detailed site characteristics.

The vegetation in the study region between 2,000 and 4,000 m a.s.l. is formed by mixed forests with dominance of *Abies pindrow, Aesculus indica, Cedrus deodara, Pinus wallichiana, Picea smithiana, Quercus semecarpifolia* and *Rhododendron arboretum*. The tree line starts at around 3,000 m a.s.l.; above the tree line herbaceous species like *Aconitum heterophyllum, Iris kemaonensis, Lindelofia macrostyla, Meconopsis aculeata, Polygonum amplexicaule, Potentilla argyrophylla* and *Rumex nepalensis* dominate the grasslands (Ahmad et al., 2021). The nival zone with sparse and scattered vegetation starts at about 4,300 m a.s.l.. The lower sites are easily accessible and thus experience high anthropogenic pressure in form of grazing, tourism, and logging (Ahmad, 2021). In the alpine vegetation belt, land-use is limited to vertical transhumance of comparatively small flocks of sheep and goat.

In 2015, we established 21 permanent plots, each of 20 × 20 m in size at every 100 m increase in elevation ranging from 2,000 to 4,000 m a.s.l. and thus covering the sub-temperate to alpine vegetation belt. The plant community structure and composition in each plot was assessed by systematic sampling using the quadrat method (Bellhouse, 2005). Specifically, within each plot we marked 5 sub-plots of 5×5 m (a total of 105 sub-plots across elevation) and within these sub-plots, 25 1×1 m quadrats (a total of 525 sub-plots across elevation) were marked to record presence-absence and abundance of shrubs and herbs in the community, respectively (Appendix 1). The vegetation in the plots was studied in 2018 during the peak growing season (February to November, depending on elevation).

Each site was characterized in terms of temperature, soil nutrients and water supply. We used mean soil temperature at 10-15 cm soil depth measured during the growing season (June to September) as a proxy for the temperature conditions (Scherer Körner, 2011). The soil temperatures were measured at a 30-minute interval by data loggers (Geo Precision Environment Technology, Germany) each installed in the centre of every permanent plot.

To characterise the soil conditions, soil samples were collected from the centre of every 5×5 m sub-plot at each permanent plot, i.e., 5 replicates per plot. The soil samples were air dried. Subsequently, we measured the concentration of available nitrogen, phosphorous and potassium (AN, AP, AK, respectively) as well as water saturation (WS; a proxy for soil moisture content) following the methods provide by Ahmad et al. (2020). Study sites coordinates, temperature logger code, and mean values of abiotic factors are given in Appendix 2 Table S1.

### Trait measurements

Based on the vegetation surveys, we selected the 139 most abundant insect-pollinated species (i.e., species that were present in more than three sites and with an abundance of >3%) from which flower and phenological traits were measured. We conducted regular phenological observations from February to November 2018, we collected data for a total of 10,025 visits to 1 × 1 m quadrats during 401 visits of the 21 permanent plots during the entire study (Appendix 2; Table S2). We recorded floral abundance of each species (if present) in 25 (1 × 1 m) sub-plots on each 20 × 20 m plot. We recorded the flowering stages (bud, open flower, or senescent flower) for every individual of a species in 1 × 1 m quadrats. Within these 1 × 1 m quadrats, the percent (%) flowering of each individual (of a species) was estimated. The flowering (%) recorded for each individual of a species in 25 (1 × 1 m) quadrats were averaged to represent flowering (%) of that species’ population, for that particular permanent plot (20 × 20 m). The day of the year (DOY 1 = 1^st^ of January), on which the plant species reached 20 and 90% of floral abundance was considered as the onset, and end of flowering, respectively as described in Valencia et al. (2016) and Ahmad et al. (2021). We also estimated the total flowering duration (TFD) calculated as time span between the end and the onset of flowering (Crimmins et al., 2013). Additionally, two floral traits namely floral display area (FDA, mm^2^) and specific flower area (SFA, mm^2^ mg^-1^) were measured following the same protocols as developed for specific leaf area described by Pérez-Harguindeguy et al. (2016). The FDA is the total area of all flowers or inflorescences of one individual at the peak of flowering and is an indicator for the investment of a plant into attracting pollinators (higher values mean higher visibility, pollinator visitation rates and potentially higher fertilization success) (Fabbro & Körner, 2004; Herrera, 2009; Zhao et al., 2016). SFA is the ratio between one-sided surface area of fresh flower divided by its dry weight and flower/inflorescence biomass, a trait that represents resource investment in reproductive parts (Pérez-Harguindeguy et al., 2016). For the FDA and SFA measurements, 10 to 20 healthy, fully flowering plants were selected for each species at each plot, dug out completely, wrapped in moist towels and transported to the laboratory. Fresh flower area was measured with the help of a digital Vernier calliper (Fabbro & Körner, 2004). Collected plant species were separated into root, shoot and flowers. These were then dried in oven at 70°C for 48 h followed by weighing using a balance with a precision of 0.1 mg. Flower/inflorescences number was recorded manually for each plant species at each plot. Floral investment was calculated as:

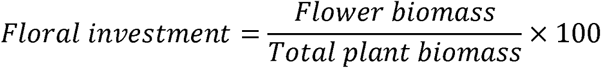

### Statistical analysis

The changes in environmental characteristics along the elevational gradient, as well as among all environmental factors, were analysed using the Pearson correlation coefficient. Polynomial regressions were applied to test the effect of rainfall and temperature during the sampling period. A detrended correspondence analysis (DCA) was performed to analyse the relationship between species occurrence and environmental parameters by using the *vegan* package in R statistical environment (Oksanen et al., 2015). An environmental fitting test with 999 permutations was performed to analyse correlations of environmental factors with the DCA axes.

The changes in community functional structure were analysed with the help of community weighted means (CWMs) and functional diversity (FD). A CWM is the average trait value weighted by species’ relative abundances in a community and thus describes the adaptation strategy of dominant species to given environmental conditions (de Bello et al., 2021). The FD is a measure of trait dissimilarity compared to random expectation (i.e., trait convergence or divergence) and was calculated using Rao’s quadratic entropy (Rao, 1982). The deviation of observed FD from the null expectation, i.e., communities are a random selection of species from the regional pool was tested with the help of null models. We generated null models by permuting the columns (species) of the plot-by-species abundance matrix for each plot (community). Then the standard effect sizes (SES) were calculated to evaluate the deviations of observed functional diversity values from random expectations. SES were calculated as the observed FD value minus the mean of the null-FD divided by the standard deviation of the null-FD. Negative SES indicated that FD was convergent, i.e., lower than expected in the null expectation. Conversely, positive SES indicated that FD is divergent, i.e., higher than the null expectations. Further, we assessed the significance of SES by identifying the proportion of random values that fell below the observed diversity value. The ranks below 0.05 indicated that FD for a given site was a significant low FD, the ranks above 0.95 indicated significant high FD.

The variability of the communities (CWM and FD) along the elevational gradient was estimated with the help of a polynomial regression (i.e., community trait value ∼ elevation + elevation^2^), to account for possible non-linear trait-elevation relationship. If the quadratic term was not significant, the term was removed.

Model assumptions were checked (e.g., homogeneity of variances and normality of residuals) and met in all cases. The flower display area was log-transformed to improve the normality of the residuals. All statistical analyses were performed in R version 4.1.0 (R Core Team 2021).

## Results

### Variability of environmental conditions along elevational gradient

Soil temperature (*r*=-0.98, p<0.001) and available phosphorus in soil (*r*=-0.65, p<0.001) strongly declined with increasing elevation (Fig. 2). Soil available nitrogen was positively correlated with soil available potassium (*r*=0.59, p<0.001) and soil moisture (*r*=0.66, p<0.001). Temperature and precipitation peaked from June to September in the study area (Appendix 2; Fig. S1). The eigenvalue of the first detrended correspondence axis was 0.59 and for the second axis 0.32 (Appendix 2; Fig. S2). Elevation and temperature reflected the main axis of variation in the multivariate space. Elevation showed negative correlation with temperature, woody cover, and available phosphorus, whereas available potassium and available nitrogen showed strong positive correlation with water saturation (Appendix 2; Fig. S2). High gradient length (DCA axis 1= 4.22) suggests high β-diversity. The sites on the extreme left of the gradient (S20, S17) share very few similarities with sites on the extreme right (S5, S6) of DCA axis 1, in terms of environmental factors and vegetation composition (Appendix 2; Fig. S2).

**Figure 2.**
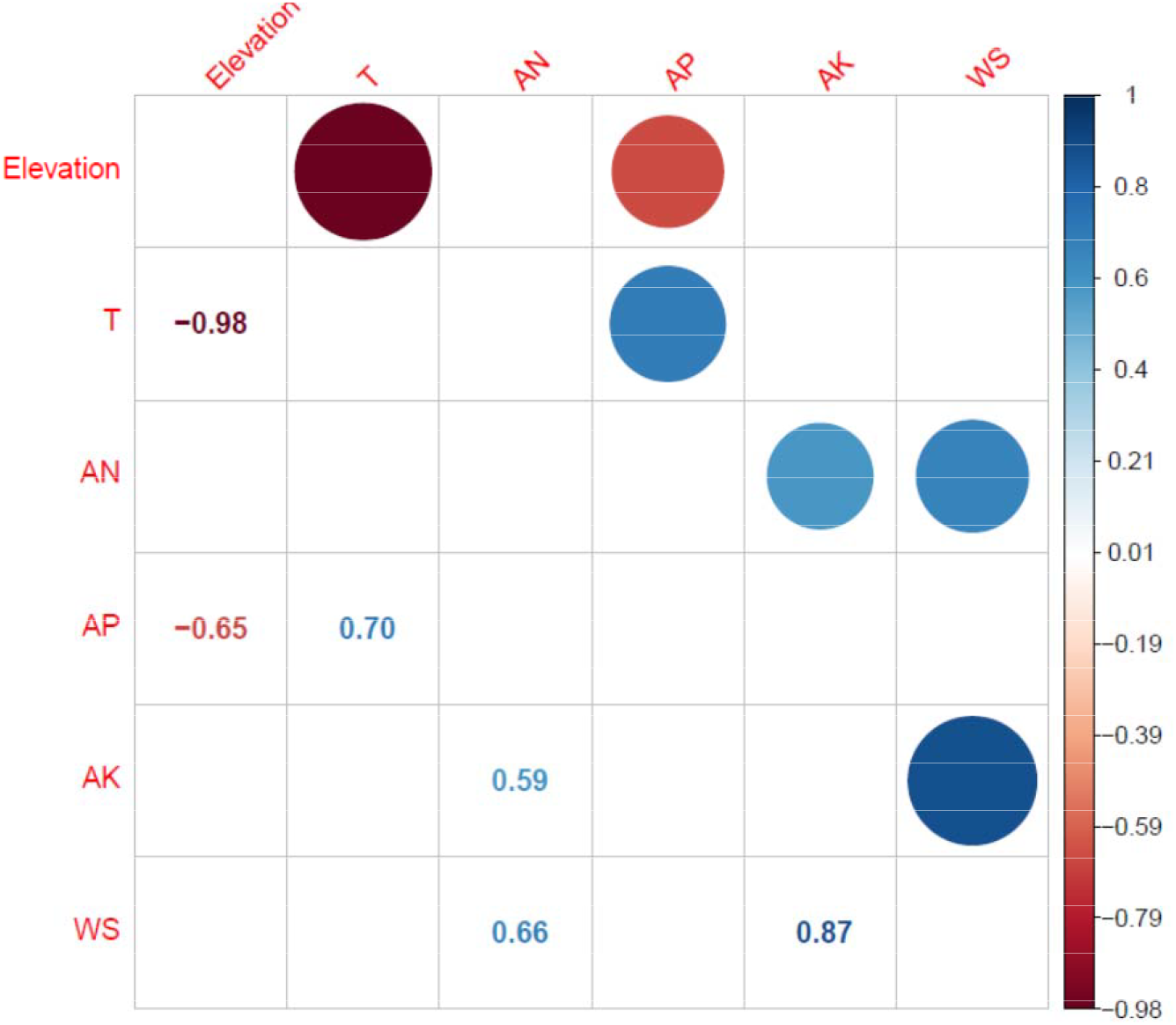
Correlation matrix for environmental factors measured in the study sites along the elevational gradient.

### Variation of community weighted means along the elevational gradient

CWMs of three out of six traits changed significantly along the elevation gradient (Fig. 3). Onset of flowering increased strongly with increasing elevations from the third week of May (DOY 140) at the lower end of the gradient to the second week of July (DOY 190) in the high elevation areas (Fig. 3A, R^2^ = 0.84, p<0.001). Specific flower area displayed an opposite pattern and decreased significantly from 19.6 mm^2^ mg^-1^ at 2,100 m a.s.l. (S2) to 9.2 mm^2^ mg^-1^ at 3,800 m a.s.l. (S19 site; Fig. 3D, R^2^ = 0.45, p<0.001). Floral display area increased with increasing elevation in a non-linear manner (Fig. 3E, R^2^ = 0.69, p<0.001) ranging from 8,302 mm^2^ at 2,500 m (S6) to the 13,7967 mm^2^ at 3,100 m (S10) a.s.l.. In ecological terms, these patterns suggest that dominant species at high elevations tended to flower later, had larger display areas, and were composed of flowers with comparatively more resource-intensive flowers (i.e., lower SFA values). The duration of flowering, flower number and flower investment did not show any significant variation with elevation (Fig. 3B, C and F, respectively).

**Figure 3.**
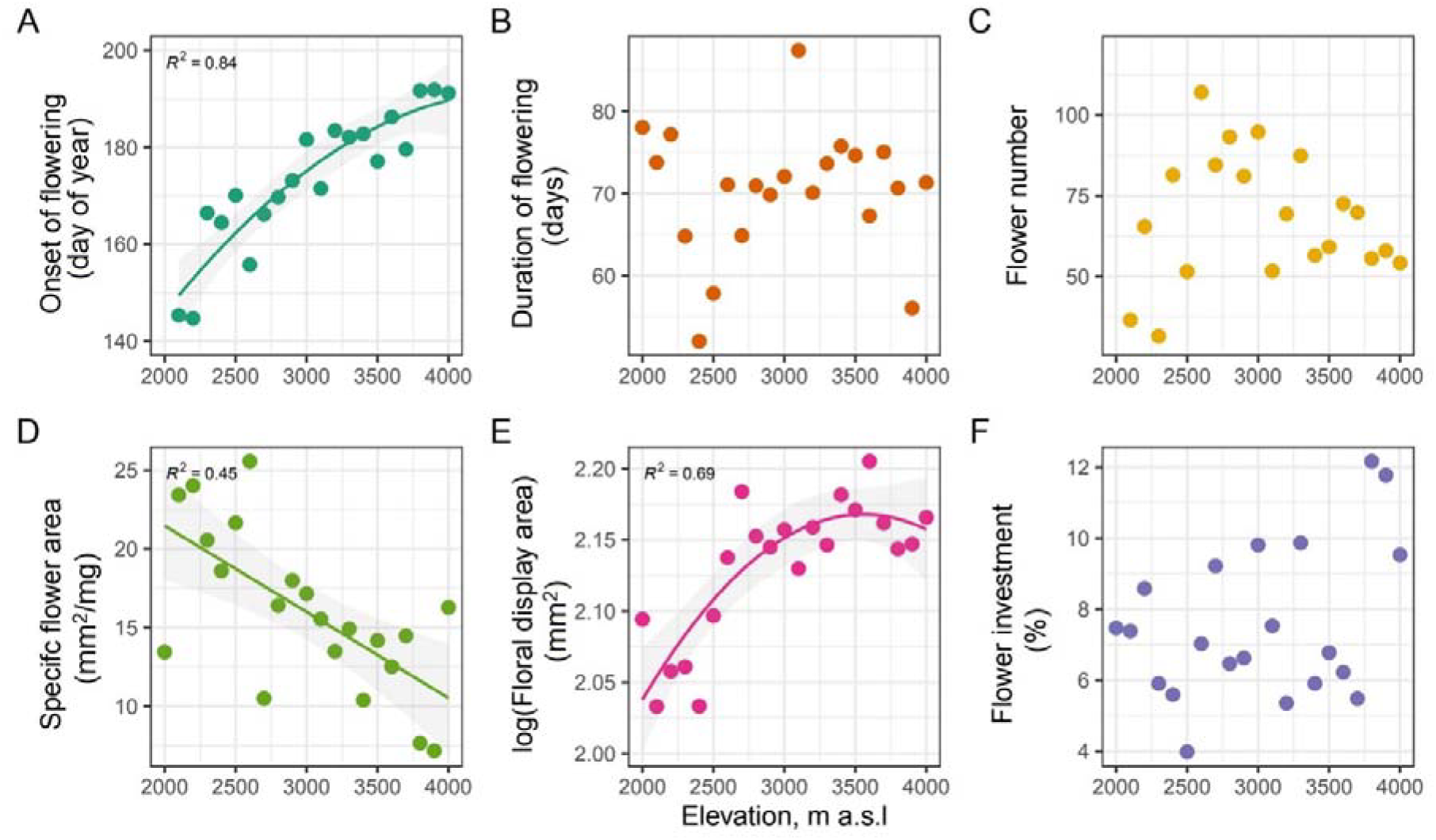
Variation of community weighted means of six traits along the elevational gradient. Lines indicate significant relationships (p<0.05), the model error is given as a shading.

### Variation of functional diversities along the elevational gradient

The FDs of almost all floral traits were non-random across the elevations (Fig. 4). FD of flowering onset decreased strongly from being divergent at lower elevations to comparatively strong trait convergence in high elevation areas (Fig. 4A; R^2^ = 0.83, p<0.001). Additionally, the FD for specific flower area displayed a significant decrease along the gradient towards more convergent trait values at higher elevations, yet the amount of variance explained by this relationship was moderate (R^2^ = 0.32, p=0.007). The FD of the remaining traits did not show any significant relationship with the elevation (Fig. 4).

**Figure 4.**
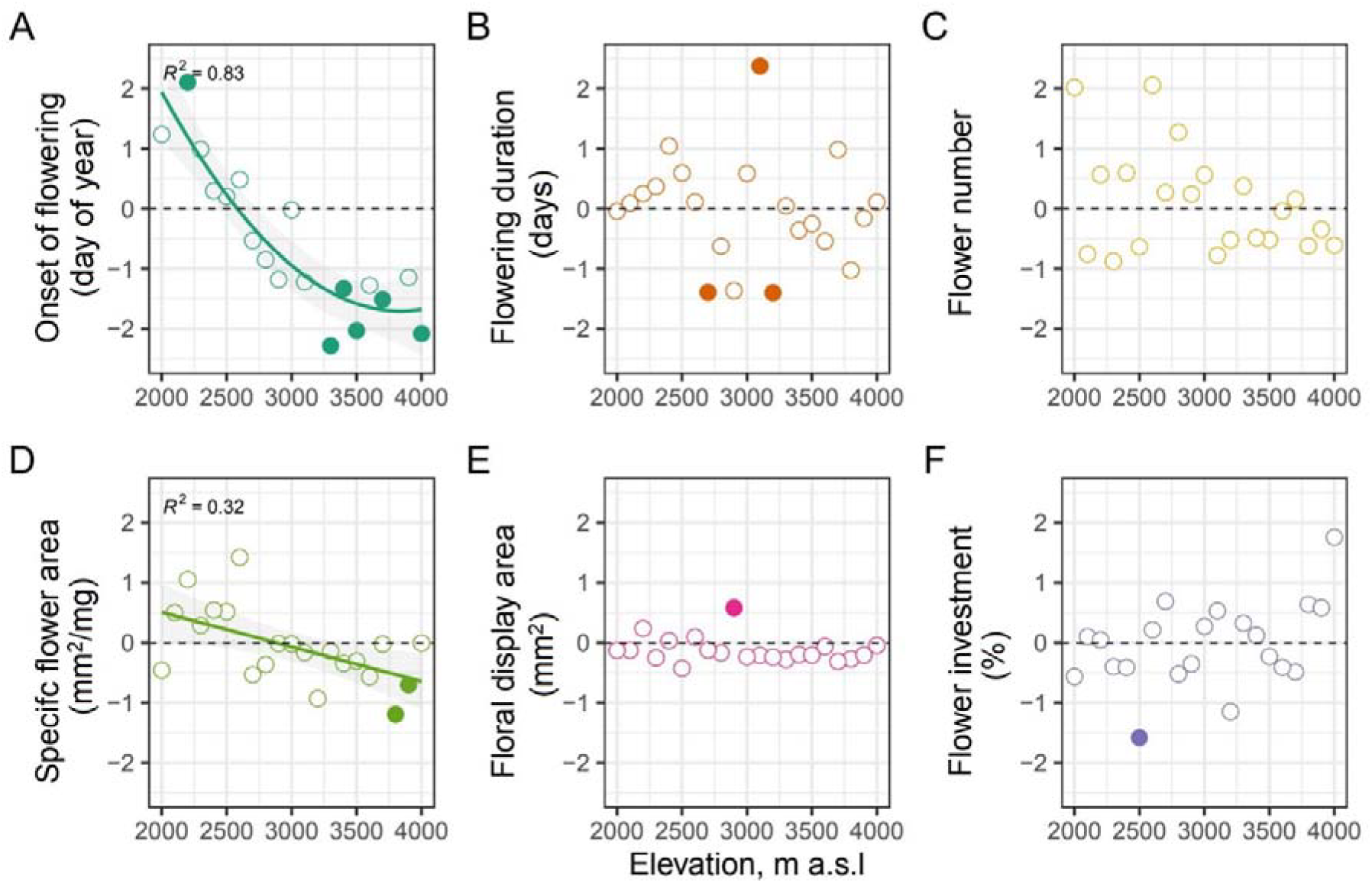
Variation of functional trait diversity (FD) of six traits with elevation. Lines indicate significant relationships according to the linear regressions (p<0.05). Negative and positive standard effect size (SES) indicate a narrower and broader trait range that expected, respectively. Filled circles represent significantly low (rank lower than 0.05) or high (rank above 0.95) FD.

## Discussion

Environmental effects on plant regeneration have long been considered as a key to understand plant population dynamics and species distribution patterns (Grubb, 1977; Woodward & Woodward, 1987), yet they have been largely neglected in community ecology research (Jiménez□Alfaro et al., 2016; Poschlod et al., 2013; Saatkamp et al., 2019). Particularly, floral traits, the main determinants of successful sexual reproduction, are rarely considered in research on community assembly (E-Vojtko et al., 2020). In this study, we close this gap and demonstrate that the so far underused phenological and morphological floral traits are important for plant community structure, especially in high elevational sites. Specificially, we reveal that abiotic filters affect phenological traits (e.g. onset of flowering), whereas plant-pollinator interactions act on flower morphological traits (e.g. floral display area) at the community level.

As expected, the temperature decreased along the elevational gradient (Fig. 2) and was the main factor determining the vegetational composition (DCA; Appendix 2 Fig. S2). The low temperature condition selected for late-flowering species (significantly higher CWM values for onset of flowering at higher elevations; Fig. 3). The strong filtering effects of low-temperature could be seen further in the convergent functional diversity of SFA in these sites (Fig. 4A & D) suggesting exclusion of maladapted early-flowering species. From the ecological point of view, these patterns can be interpreted as a plant phenological strategy to avoid frost injury in frost-susceptible flowers (Wagner et al., 2011), which seem to be a general adaptation of plant reproduction to alpine environments (Körner, 2021). Alternatively, the delayed and synchronised onset of flowering in high elevation areas (strong trait convergence suggest a comparatively low number of available temporal niches for flowering) could be adaptive to facilitate pollination success under conditions with low pollinator abundancy (‘magnet’ effect; Koski & Ashman, 2015; Bucher & Römermann, 2020). Further, delayed flowering with increasing elevation might facilitate plant-pollinator phenological overlap, as pollinator activity at the higher elevation peaks in the second half of the vegetation period, when temperatures are high enough (Rawat, 2012). In contrast, the lower elevational sites seem to harbour a large diversity of plants with different phenological strategies aiming at reducing plant-plant competition for pollinators (Zhao et al., 2016).

Interestingly, the significantly later onset of flowering did not affect the duration of flowering in high elevation communities both expressed as CWMs and FDs (Fig. 3B and 4B). These findings contradict previous observations made in other regions that flowering duration decreases with increasing elevations, due to decreasing time window for successful pollination, fertilization, and consequently seed production (Bucher & Römermann, 2020; Körner, 2021). The most plausible explanation is that the onset of monsoon season in the study system, which coincides with the peak of flowering, i.e., the middle of the growing period in the Himalayan alpine vegetation belt (Ram et al., 1988), may relax the low-temperature filtering effects on plant phenology and thereby promoting longer duration of flowering. The comparatively long flowering in the resource-limited harsh alpine environments might be achieved by considerably longer life spans of cost-effective flowers (Trunschke & Stöcklin, 2017; see below). Cumulatively, these findings contradict our hypothesis and suggest that flowering duration plays no important role in the community assembly process in the study system.

Analyzing morphological floral traits, we found that CWMs for specific flower area and floral display significantly varied along the elevation gradient with high elevation communities dominated by species with large flower display areas and relatively small SFA values (Fig. 3D). Most likely, these shifts in CWM can be related to the effects of environmental filtering and plant-pollinator interactions on sexual regeneration strategies in alpine environments. As for the former, we assume that low-temperature stress and lower soil fertility (Fig. 2) as well as stronger winds (Körner, 2021) at higher elevational sites select for plants that invest more resources per floral display area unit. In alpine regions, flowers adapt in response to wind induced mechanical perturbations, as heavier flowers have more mechanical tissue that supports the floral unit, making the structures more flexible and aerodynamically stable (Cordero et al., 2007; Zhang et al., 2021). The larger flower display area might be also adaptive in cold alpine climates as it positively correlated with a flowers’ passive heat accumulation (Dietrich and Körner, 2014). As for the plant-pollinator interactions, the decreased SFA CWM and FD values can be also attributed to the increased longevity of single flowers at higher elevations to maximize pollination success under conditions of low pollinator abundancy (Fabbro & Körner 2004; Klatt et al., 2018). Similarly, a larger display area allows plants from alpine communities to increase pollination probability by attracting more pollinators (Fabbro & Körner 2004; Kiełtyk, 2021; Zhao et al., 2016). Regardless of the main driver of SFA variability at the community level, this trait seems to play an important role in community assembly process as we observed a significantly higher trait convergence with increasing elevation.

Contrastingly to previous studies in alpine environments (Fabbro & Körner, 2004), resource investment in reproduction (‘flower investment’) did not change along the elevational gradient. Many studies suggested increase in pollinator visitation rates with increase in flower size and number (Conner & Rush, 1996). However, a reason for no significant change in flower number with elevation might be that with increasing flower display size the plant does not need to invest more resources to flower number. In addition, Samson and Werk (1986) asserted that for understanding the pattern of resource investment in plants, the species-specific size-dependent effects should be taken into consideration. Obeso (2002) reported that the responses to plant reproductive investment are extremely variable due to previous history, individual variations, timing of reproduction, plant size, plant density, competition, plant architecture, and successional stage.

Thus, the interaction of different life-history traits and plant size effects with reproductive investment should be studied to understand the resource allocation strategy of plants across elevation.

## Conclusions

The revealed patterns of CMW and FD variations of floral traits have at least three important points for plant ecological research. First, our study clearly demonstrates that in addition to widely used vegetation traits, floral traits can also play a significant role in community assembly process along environmental gradient. Therefore, these finding provide further support to the recent argument that plant research should consider multiple traits representing different ecological functions including growth, reproduction, and survival (E-Vojtkó et al. 2020; Laughlin, 2014; Rosbakh et al., 2021).

Second, our findings suggest that the floral traits might be used in applied ecological research to forecast functional community composition shifts in alpine ecosystems. The continuous temperature rise observed in the study region (Sabin et al., 2020) may lead to relaxation of low-temperature stress filtering effects on flowering phenology, specific flower area and floral display area of species adapted to environmental conditions of lower sites, resulting in an altered composition of communities in cold sites (i.e., increase in frequency and abundance of species from lower elevations). The results also revealed a lacuna in our understanding of reproductive investment patterns of species and its interactions with biotic and abiotic constraints along elevation, which might provide additional insights into the factors governing community assembly of flowering plants.

## Author contributions

MA, SR, SFB, HPS, DRB and SKU conceive the idea and design the study for this work. MA, SRA, PS and SKU collected the data. MA, SRA, and PS data entry and did the formal analyses. SR and MA analysed the data. MA and SR wrote the first version of the manuscript; all authors contributed to writing of the final version of the manuscript.

## Acknowledgements

Authors are grateful to Ministry of Environment, Forest & Climate Change, India, for providing financial support under National Mission on Himalayan Studies. MA is thankful to Council of Scientific and Industrial Research (CSIR), India, for providing senior research fellowship. Authors express gratitude to Sanjay Kumar (Director, CSIR-IHBT) who provided the mandatory facilities for the research work at CSIR-IHBT. We also acknowledge Christian Körner and Robert R. Junker for insightful suggestions during the initial stages of project proposal. We acknowledge Rohit, Om Prakash, Girja Nand, and Ashok Kumar for their assistance in carrying out the fieldwork.

## Data accessibility statement

Should be manuscript be accepted for publication, the data will be published on Zenodo.

## Notes

### Competing Interest Statement

The authors have declared no competing interest.

### Summary of Updates

Supplemental file updated

