## Supplementary material for "The role of floral traits in community assembly process at high elevations in Lesser Himalaya": Table S1

**Appendix 2**

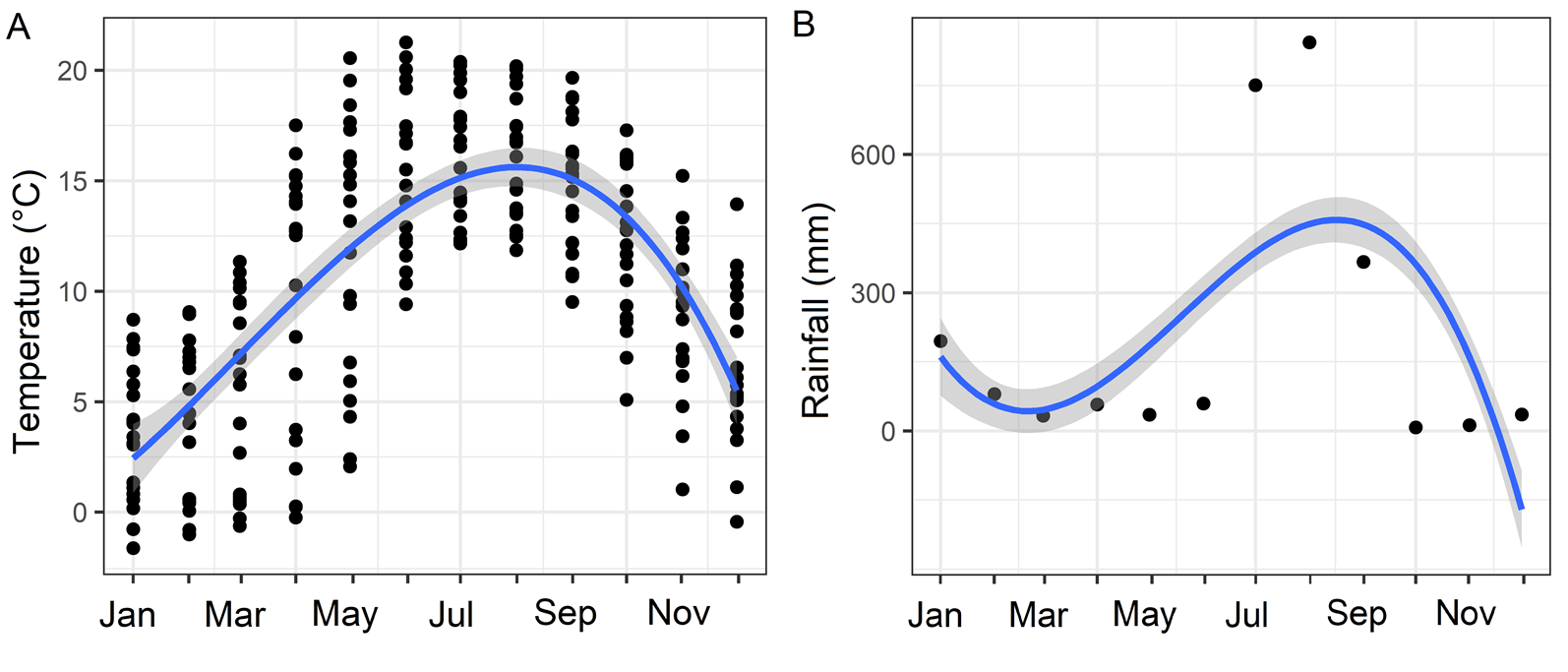

**Figure S1:** Monthly pattern of temperature and rainfall in the study area. A) temperature and B) rainfall pattern during sampling period.

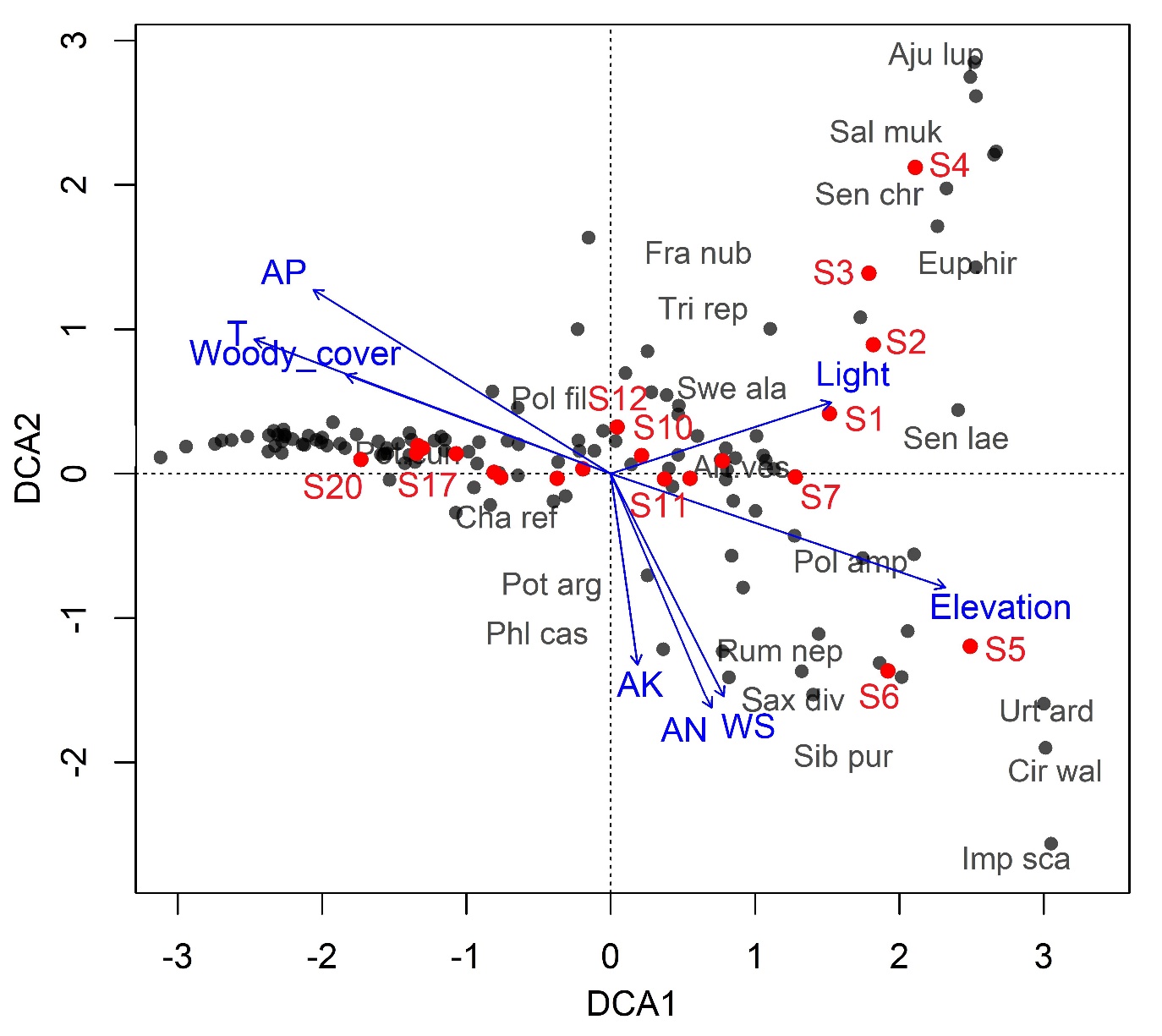

**Figure S2:** Plot of the first two axes of a Detrended Correspondence Analysis (DCA) on species composition, explaining 74.79 % of the total variance. ‘S’ represents the different elevational sites. Arrows depict correlations of environmental factors with the DCA from an environmental fitting analysis. Abbreviations: AN = available nitrogen; AK = available potassium, AP = available phosphorus; T = temperature and WS = water saturation. Species names are printed in black and sites are red.

**Table S1:** Study sites description, temperature logger code, coordinates along with mean values of abiotic factors.

| **Sites** | **Temperature logger code** | **Latitude(N)** | **Longitude(E)** | **Elevation (m a.s.l.)** | **T**  **(°C)** | **Light intensity (μmol m^−2^S^−1^)** | **AN**  **(kg/ha)** | **AP**  **(kg/ha)** | **AK**  **(kg/ha)** | **WS (%)** | **Woody_cover (%)** |
| --- | --- | --- | --- | --- | --- | --- | --- | --- | --- | --- | --- |
| S1 | A52033 | 32°08'19.813'' | 76°33'47.856'' | 2000 | 15.34771 | 1525.667 | 357.1133 | 7.176667 | 170.0133 | 48.79 | 51.05 |
| S2 | A52041 | 32°08'28.330'' | 76°33'35.495'' | 2100 | 14.19033 | 1436.095 | 209.1433 | 18.56667 | 166.1433 | 49.48333 | 40.86194462 |
| S3 | A52037 | 32°08'39.081'' | 76°33'40.890'' | 2200 | 13.09486 | 1444.524 | 196.2633 | 7.113333 | 174.1767 | 43.59333 | 40.97 |
| S4 | A5139F | 32°08'48.127'' | 76°33'43.619'' | 2300 | 14.37058 | 1444.238 | 379.44 | 10.84 | 206.8933 | 52.26667 | 60.76260897 |
| S5 | A5203B | 32°09'16.076'' | 76°34'13.482'' | 2400 | 14.29344 | 1539.238 | 305.8267 | 25.38667 | 270.8133 | 50.59 | 7.37127887 |
| S6 | A52035 | 32°09'18.972'' | 76°34'21.624'' | 2500 | 13.64802 | 1501.571 | 306.23 | 30.68 | 278.8967 | 53.35333 | 10.5985854 |
| S7 | A52039 | 32°09'19.549'' | 76°34'30.359'' | 2600 | 12.47098 | 1563.048 | 361.0133 | 15.24333 | 289.4233 | 56.24667 | 30.64119582 |
| S8 | A5203C | 32°09'22.290'' | 76°34'35.450'' | 2700 | 11.47529 | 1576.429 | 279.4167 | 11.34 | 272.51 | 54.25333 | 16.59 |
| S9 | A5203D | 32°09'25.413'' | 76°34'37.458'' | 2800 | 11.86958 | 1457.333 | 297.3467 | 6.89 | 290.5933 | 59.26667 | 6.1 |
| S10 | A52038 | 32°09'36.088'' | 76°34'38.665'' | 2900 | 11.0395 | 1561.429 | 369.65 | 13.95333 | 320.92 | 58.72 | 4.22 |
| S11 | A52034 | 32°09'43.349'' | 76°34'39.421'' | 3000 | 10.92378 | 1602.238 | 292.31 | 6.646667 | 261.0367 | 53.94333 | 1.77 |
| S12 | A52040 | 32°09'53.379'' | 76°34'38.695'' | 3100 | 9.875626 | 1590.778 | 316.6633 | 12.56667 | 294.08 | 55.67333 | 0 |
| S13 | A5203A | 32°09'59.572'' | 76°34'45.963'' | 3200 | 8.672457 | 1580.444 | 350.0433 | 6.36 | 303.35 | 59.42333 | 0 |
| S14 | A51396 | 32°10'05.820'' | 76°34'55.776'' | 3300 | 8.52 | 1554.389 | 397.48 | 4.693333 | 298.1 | 56.70667 | 2.25 |
| S15 | A52036 | 32°10'10.083'' | 76°36'10.479'' | 3400 | 7.039649 | 1606 | 369.5733 | 10.35333 | 292.3433 | 55.52 | 6.85 |
| S16 | NA | 32°11'02.532" | 76°36'02.281" | 3500 | 7.496146 | 1675.214 | 361.99 | 6.236667 | 227.44 | 53.64333 | 9.52 |
| S17 | A5203E | 32°11'02.730" | 76°36'02.505" | 3600 | 6.687024 | 1694.071 | 339.3333 | 2.524 | 222.3633 | 52.49467 | 2.263420354 |
| S18 | A5203F | 32°11'12.654" | 76°36'04.118" | 3700 | 5.906399 | 1616.429 | 149.87 | 0.786667 | 192.6933 | 44.04333 | 0 |
| S19 | A52030 | 32°11'15.833" | 76°36'01.858" | 3800 | 6.456756 | 1670.429 | 152.7367 | 0.68 | 192.09 | 51.65 | 0 |
| S20 | A52042 | 32°11'27.538" | 76°36'09.786" | 3900 | 5.038794 | 1783.5 | 159.19 | 0.72 | 187.6 | 46.36667 | 0 |
| S21 | A5203 | 32°11'34.109" | 76°36'15.468" | 4000 | 4.276928 | 1757.25 | 277.7767 | 0.566667 | 157.6667 | 44.22667 | 0 |

*Abbreviations:* S1= 2000 m, S2= 2100 m, S3= 2200 m, S4= 2300 m, S5= 2400 m, S6= 2500 m, S7= 2600 m, S8= 2700 m, S9= 2800 m, S10= 2900 m, S11= 3000 m, S12= 3100 m, S13= 3200 m, S14= 3300 m, S15= 3400 m, S16= 3500 m, S17= 3600 m, S18= 3700 m, S19= 3800 m, S20= 3900 m, S21= 4000 m. T= temperature, AN= available nitrogen, AP= available phosphorus, AK= available potassium, WS= water saturation.

**Table S2:** Date, elevational plots survey and flower phenology observation.

| **Month** | **Date** | **Elevation** | **Total sites** | **Observational Quadrats at each site (1x1 m^2^)** | **Total observational quadrats (1x1 m^2^)** |
| --- | --- | --- | --- | --- | --- |
| February | 10 | S1,S2 | 2 | 25 | 50 |
| February | 19 | S1,S2,S3 | 3 | 25 | 75 |
| February | 28 | S1,S2,S3,S4 | 4 | 25 | 100 |
| March | 10 | S1,S2,S3,S4 | 4 | 25 | 100 |
| March | 20 | S1,S2,S3,S4,S5 | 5 | 25 | 125 |
| March | 30 | S1,S2,S3,S4,S5 | 5 | 25 | 125 |
| April | 9 | S1,S2,S3,S4,S5 | 5 | 25 | 125 |
| April | 19 | S1,S2,S3,S4,S5,S6 | 6 | 25 | 150 |
| April | 29 | S1,S2,S3,S4,S5,S6 | 6 | 25 | 150 |
| May | 10 | S1,S2,S3,S4,S5,S6,S7 | 7 | 25 | 175 |
| May | 20 | S1,S2,S3,S4,S5,S6,S7,S8 | 8 | 25 | 200 |
| May | 30 | S1,S2,S3,S4,S5,S6,S7,S8,S9 | 9 | 25 | 225 |
| June | 10 | S1,S2,S3,S4,S5,S6,S7,S8,S9,S10,S11 | 11 | 25 | 275 |
| June | 19 | S1,S2,S3,S4,S5,S6,S7,S8,S9,S10,S11,S12,S13 | 13 | 25 | 325 |
| June | 30 | S1,S2,S3,S4,S5,S6,S7,S8,S9,S10,S11,S12,S13,S14,S15 | 14 | 25 | 350 |
| July | 10 | S1,S2,S3,S4,S5,S6,S7,S8,S9,S10,S11,S12,S13,S14,S15,S16,S17,S18 | 18 | 25 | 450 |
| July | 18 | All | 21 | 25 | 525 |
| July | 29 | All | 21 | 25 | 525 |
| August | 10 | All | 21 | 25 | 525 |
| August | 20 | All | 21 | 25 | 525 |
| August | 30 | All | 21 | 25 | 525 |
| September | 10 | All | 21 | 25 | 525 |
| September | 20 | All | 21 | 25 | 525 |
| September | 30 | All | 21 | 25 | 525 |
| October | 10 | All | 21 | 25 | 525 |
| October | 20 | All | 21 | 25 | 525 |
| October | 30 | All | 21 | 25 | 525 |
| November | 10 | All | 21 | 25 | 525 |
| November | 20 | S1,S2,S3,S4,S5,S6,S7,S8,S9,S10,S11,S12,S13,S14,S15 | 15 | 25 | 375 |
| November | 30 | S1,S2,S3,S4,S5,S6,S7,S8,S9,S10,S11,S12,S13 | 14 | 25 | 350 |
| Total |  |  | 401 |  | 10025 |

*Abbreviations:* S1= 2000 m, S2= 2100 m, S3= 2200 m, S4= 2300 m, S5= 2400 m, S6= 2500 m, S7= 2600 m, S8= 2700 m, S9= 2800 m, S10= 2900 m, S11= 3000 m, S12= 3100 m, S13= 3200 m, S14= 3300 m, S15= 3400 m, S16= 3500 m, S17= 3600 m, S18= 3700 m, S19= 3800 m, S20= 3900 m, S21= 4000 m.
